# Using wearable biosensors and ecological momentary assessments for the detection of prolonged stress in real life

**DOI:** 10.1101/2021.06.29.450360

**Authors:** Rayyan Tutunji, Nikos Kogias, Bob Kapteijns, Martin Krentz, Florian Krause, Eliana Vassena, Erno J. Hermans

**Author notes:** **Corresponding Author**, Rayyan Tutunji.

## Abstract

**Background:** Increasing efforts toward prevention of stress-related mental disorders have created a need for unobtrusive real-life monitoring of stress-related symptoms. Wearable devices have emerged as a possible solution to aid in this process, but their use in real-life stress detection has not been systematically investigated.

**Methods:** Using ecological momentary assessments (EMA) combined with wearable biosensors for ecological physiological assessments (EPA), we investigated the impact of an ecological stressor (i.e., an exam week) on physiological arousal and affect. With this paradigm we investigated whether we could use wearable devices to detect stress states using machine learning models.

**Results:** During stressful high-stake exam (versus control) weeks, participants reported increased negative affect and decreased positive affect. Intriguingly, physiological arousal was decreased on average during the exam week. Time-resolved analyses revealed peaks in physiological arousal associated with both self-reported stress and self-reported positive affect, while the overall decrease in physiological arousal was mediated by lower positive affect during the stress period. We then used machine learning to show that a combination of EMA and physiology yields optimal identification of stress states.

**Conclusions:** Our findings highlight the potential of wearable biosensors in stress-related mental-health monitoring, but critically show that psychological context is essential for interpreting physiological arousal detected using these devices.

## Introduction

Stress-related mental disorders such as major depression and anxiety disorders have gained increased recognition in the public eye. While a vast body of research exists regarding these disorders, studies have mostly focused on retrospective assessments of afflicted individuals. More recently, an increased interest has emerged in determining what makes some individuals more resilient to developing these disorders than others(Hermans & Fernández, 2015; Kalisch et al., 2017; McEwen, 2016; Osório et al., 2017). Investigating resilience requires investigation of individual variation in stress reactivity prior to the development of psychological illness(Kalisch et al., 2017). A driving force behind this approach is the need to establish early warning signs of subsequent onset of stress-related disorders. Early interventions improve psychological outcomes in patients(Giummarra et al., 2018), and reduce societal and economic burdens of psychiatric illness on society(Reynolds et al., 2012). The ability to unobtrusively detect states of stress in daily life would enable early ecological interventions in those at risk by either flagging risk states to health-care providers, or by delivering in-the-moment personalized interventions during these periods(McDevitt-Murphy et al., 2018).

Previous studies have used Ecological Momentary Assessments (EMA)(Shiffman et al., 2008) to derive ecologically valid experiences of stress in daily life. These paradigms use repeated questionnaires (“beeps”) in daily life to investigate various psychological processes(Bar-Kalifa & Sened, 2019; Collip et al., 2013; Swendsen, 2016). Such methods used in stress-related disorders have identified real-life behavioral patterns that may explain or predict onset of psychiatric illness(Wichers et al., 2019). They have also given insight into effects of stress exposure on mood and its links to depression(Dunkley et al., 2017). Despite providing substantial insights, these methods are often intrusive (i.e. require active participation of patients), can lack feasibility in psychiatric populations, and may be influenced by careless responses, or lower subjective insight into symptoms and associated states(Quee et al., 2011). Furthermore, the sparse sampling of subjective states may miss time windows in which stressors occur. These issues indicate a growing need for novel and more reliable methods for passive and ambulatory mental-health monitoring.

The emergence of widely accessible wearable biosensors has raised the question whether these devices can be used for ecological *physiological* assessments (EPA), either as an add-on or an alternative to EMA, in mental health monitoring. Wearable biosensors offer continuous recording of autonomic physiological markers such as skin conductance (SC) and heart rate (HR). These measures have been extensively validated in laboratory-based studies using controlled stress-induction protocols(Schwabe et al., 2008), showing increased HR and SC and decreased HR variability in response to stressors(Löw et al., 2008; Pereira et al., 2017). However, these autonomic physiological parameters are also associated with general arousal(Lang et al., 1993), including high-arousal states for positive affect(Löw et al., 2008). Thus, using EPA may be more complicated in daily life than in the lab: While acute stress may trigger arousal, arousal itself may not necessarily signal the presence of stress. The relationship of autonomic physiological responses to stressors in real life is not well understood.

Some studies have attempted to investigate the physiology of daily life stress, using scenarios or methods that are restrictive or burdensome(Healey & Picard, 2005; Hovsepian et al., 2015). For instance, a study using wearable biosensors could replicate lab findings to some extent(Smets et al., 2018). However, this study lacked an environmental stressor and relied on the assumption that subjective stress measures can be taken as the “ground truth”. Overall reports of stressed states in this study were also relatively low when compared to the non-stress states. Finally, it did not allow probing the consequences of accumulation of stress over a prolonged period, a key aspect when considering mental health.

To this end, we aimed to investigate the validity of passive EPA monitoring of physiological arousal and active EMA measures to detect prolonged stress exposure. We investigated a population of first-year medical and biomedical students, known to experience increased psychological distress(Maser et al., 2019). Participants collected EMA and EPA data once during a week culminating in a high-stake examination (i.e., stress week) and another without (i.e., control week). We first validated our protocol by testing between-week differences in EMA subjective stress. We then assessed the impact of exam/stress weeks on mood and physiology outcomes. Finally, we used individualized machine-learning models to classify per time point (beep) which week participants were in using either mood, physiological, or a combination of both measures. We predicted increased autonomic physiological responses and negative affect, and decreased positive affect. We expected that both EPA and EMA measures would successfully identify prolonged stress states, and predicted that models combining EPA and EMA would outperform the single models.

## Materials and Methods

### Experimental Design

We recruited 84 right-handed, first year bachelor’s students in the medical or biomedical science majors from Radboud Health Academy spanning three academic years (2017, 2018, and 2019). One participant withdrew during testing, resulting in a total sample size of 83 participants used in the analysis. The programs were selected due to their structured examination weeks that occur every 5^th^ and 10^th^ week of a semester, allowing us to examine a period with higher stress levels during examination weeks as an ecological prolonged stressor. Only participants with no history of psychiatric illness were included in the study. Recruitment was stopped following the COVID-19 outbreak (March 2020). All procedures carried out were approved by the regional medical ethical review board (CMO Arnhem-Nijmegen). The authors assert that all procedures contributing to this work comply with the ethical standards of the relevant national and institutional committees on human experimentation and with the Helsinki Declaration of 1975, as revised in 2008.

Participants completed two weeks of ecological momentary assessments (EMA), one during an examination period (i.e., stress week) and the other occurring on average 16 days (min=10, max=33) outside of these periods (i.e., control week, demographics in Table 1). We maintained at least one week between the end of one week and the start of the other to ensure sufficient recovery time from the stressor. Compliance rates were overall high with 84% of surveys completed within the allocated one-hour window during both weeks. When accounting for missing and poor-quality physiology (EPA) data, completion rates dropped to between 76 −77% (within median ranges for EMA studies)(Vachon et al., 2019). Gender distribution was similar to that of students enrolled at the university (57% female, according to Radboud University website). We were unable to fully counterbalance the order of weeks due to early termination of recruitment, but instead controlled for it in all statistical analyses.

**Table 1.**
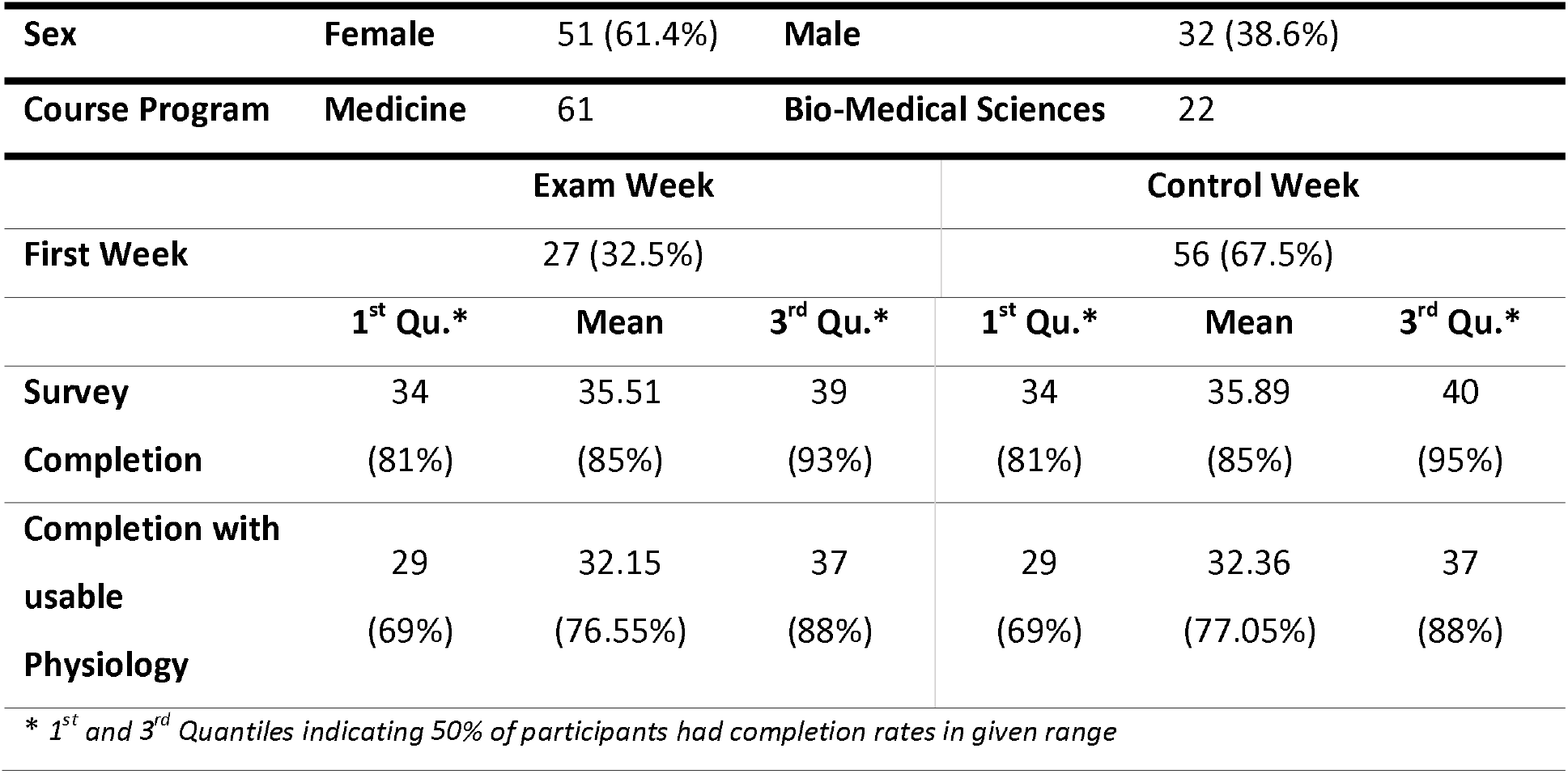
Descriptive statistics.

Participants also filled in questionnaires and participated in MRI sessions which are outside the scope of the current paper and will be reported elsewhere.

### Assessing daily-life stress through EMA and EPA

The comparison of the stress week (exam week) vs control week allowed us to determine individualized patterns of stress reactivity. During these weeks, participants received six surveys a day at fixed intervals via SMS-text links. Participants were given a one-hour window to fill in the surveys (like previous studies(Schultchen et al., 2019)). Surveys assessed different psychological aspects related to stress, including event, activity, social, and physical stress as well as positive (PA) and negative (NA) affect outcomes (see SM1 for surveys). The first questionnaire of the day contained a sleep quality assessment, and the last a self-reflection questionnaire. Participants were instructed to wear an Empatica E4 wristband (Empatica, Milano, Italy) recording ambulatory EPA data throughout both weeks (collected passively, continuously, and in the background). Participants were instructed to charge and synchronize the watch to researcher-specific accounts once a day for one hour. A detailed explanation was given to participants on E4 operation with a practice session during the intake interview. The E4 devices collected blood pulse volume, electrodermal activity, three-axis movement, and body temperature (See SM Text 2 for detailed EMA and EPA pipeline description).

### Statistical Analysis

All statistical analyses were conducted in R (R, version 3.6.1) using generalized linear mixed effects models and random forests (lmer and randomforest packages)(Kuznetsova et al., 2017; Liaw & Wiener, 2002). Initial analyses examined overall differences in subjective stress between the two weeks to establish the validity of the experimental manipulation. We then tested for the effect of an exam week on affect and physiology. We additionally tried replicating previous findings associating momentary stress with physiology and mood. Mediation analysis was then used to explain the apparent differences in the relationships between the week type and momentary analyses (see SM text 3 for a full report).

### Machine-Learning Models

One of our goals was assessing the usability of ambulatory, non-intrusive measures to determine whether someone is currently in a stressed state. To this end, random forest models were used to determine the ability to classify whether subjects’ beeps were in the stress or control week using the collected EMA mod and/or ambulatory EPA outcome data. We conceptualized mood and physiology as outcomes of stressed states based on previous findings(Dunkley et al., 2017; van der Stouwe et al., 2019). Individualized models were estimated using a Leave-One-Beep-Out (LOBO) approach at a single-subject level, where models were trained on individuals’ n-1 beep data and tested on the removed beep, repeating until all beeps had been removed. Three models were tested as follows: Model 1 tested the ability to classify week type from (momentary) PA and NA, Model 2 from EPA data, and Model 3 from the combination of both. Models were tested against a bootstrap error distribution (n=10,000), with group effects tested using paired sample t-tests against the mean subject-level bootstrap error. We tested the generalizability of the random forest models to a population level using a Leave-One-Subject-Out (LOSO) analysis in which models are were trained on N-1 participants dataset and tested on the removed participant, repeating until each participant had been removed once from the dataset. Model predictions using the LOBO were then compared to that of the LOSO method to estimate the generalizability of machine-learning models on the data.

## Results

### Examination periods are associated with increased self-reported stress

We found a significant increase in prominent stressful events (i.e., event-related stress, β=0.30, CI=[0.18,0.42], p<0.001) and current reports of stress (i.e., activity-related stress, β=0.51, CI=[0.30,0.71], p<0.001) in the exam stress versus control week. Social stress was not significantly different between the two weeks. The control items measuring physical stress also did not differ significantly between the weeks, showing that increases in subjective stress were likely due to our experimental manipulation, instead of environmental or physical changes (Figure 2A, SM Table 1 for full results). As anticipated, not all beeps in stress weeks were subjectively reported as stressful, while some beeps during the control week were subjectively rated as stressful. To quantify this, subjective stress variables for event, social, and activity stress were aggregated across both weeks. A median split was then used to estimate the percentage of incongruent self-report beeps (i.e., false positives in stress weeks and false negatives in control weeks). On average across participants, 45% of the beeps yielded self-reported stress incongruent with the week type. Machine learning models using self-reported stress assessments in a leave-one-beep (LOBO) approach to classify week type achieved similar error rates, with 43% of beeps being classified as the wrong week type.

**Fig 1.**
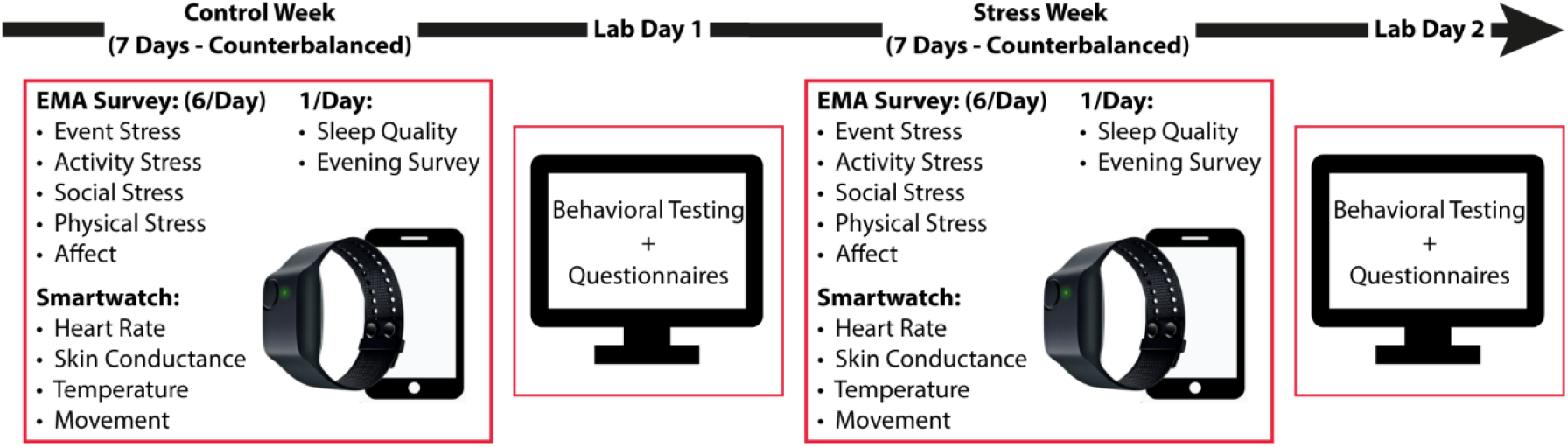
Study Timeline. Diagram portraying sequence of participation in the study with counterbalanced weeks

**Fig 2.**
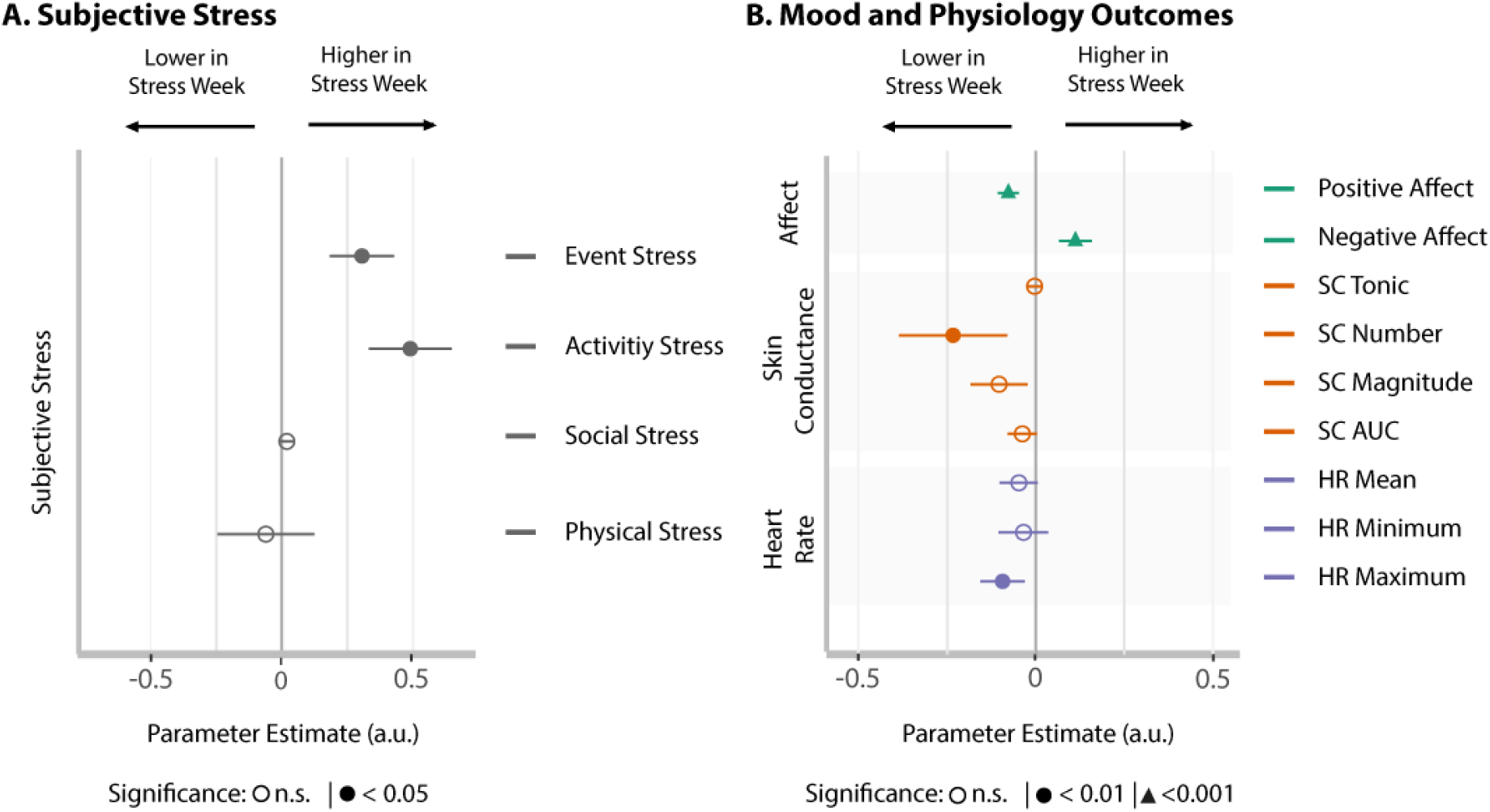
Fixed effects estimates of between-week difference. **(A)** Event-related stress (pertaining to the most prominent event since the last survey), and activity-related stress (relating to the current activity participants are engaged in) are significantly higher in the stress week compared to the control week. **(B)** This is accompanied by increased negative affect, decreased positive affect, and decreases in averages of multiple arousal-related physiological measures. Error bars represent confidence intervals.

In accordance with our expectations, we also saw an increase in negative affect (β=0.12, CI=[0.08,0.17], P_fdr_<0.001), and decrease in positive affect (β=-0.08, CI=[-0.11,-0.05], p_fdr_<0.001) during the stress week (Figure 2B). Unexpectedly, we found a decrease in physiology arousal-related measures during the exam week including the number of SC responses (log-Mean=-0.27, CI=[-0.42,-0.12], p_fdr_<0.01), and maximum HR (β=-0.10, CI=[-0.16,-0.03], p_fdr_<0.01). Figure 2B, SM Table 2 for full results).

### Momentary subjective stress is associated with mood and physiology

To explore the dynamics underlying the unexpected average decrease in measures of physiological arousal during the stress week, we investigated the link between moment-to-moment fluctuations in subjective stress and outcome measures (mood and physiological arousal). We found a positive association between negative affect and activity-related (β=0.06, CI=[0.01,0.12], p_fdr_<0.05), social (β=0.22, CI=[0.18,0.27], p<0.001), and physical stress (β=0.15, CI=[0.12,0.18], p_fdr_<0.001). The opposite was true for positive affect for event-related (β=-0.12, CI=[-0.19,-0.06], p_fdr_<0.001), activity-related (β=-0.17, CI=[-0.25,-0.09], p<0.001), social (β=-0.28, CI=[-0.34,-0.22], p_fdr_<0.001), and physical stress (β=-0.23, CI=[-0.27,-0.18], p_fdr_<0.001). The magnitude of SC responses was associated with activity (β=0.08, CI=[0.02,0.15], p_fdr_<0.05), event (β=0.07, CI=[0.02,0.13], p_fdr_<0.05) and physical stress (β=0.03, CI=[0.00,0.06], _fdr_p<0.05). For HR measures, mean (β=-0.04, CI=[0.08,-0.01], p_fdr_<0.05) and minimum (β=-0.02, CI=[-0.03,-0.00], P_fdr_<0.05) HR were negatively associated with social stress. Thus, moment-to-moment fluctuations in subjective stress are associated with expected mood changes and increases in physiological arousal (Fig 3, see SM Table 3 for full results).

**Fig 3.**
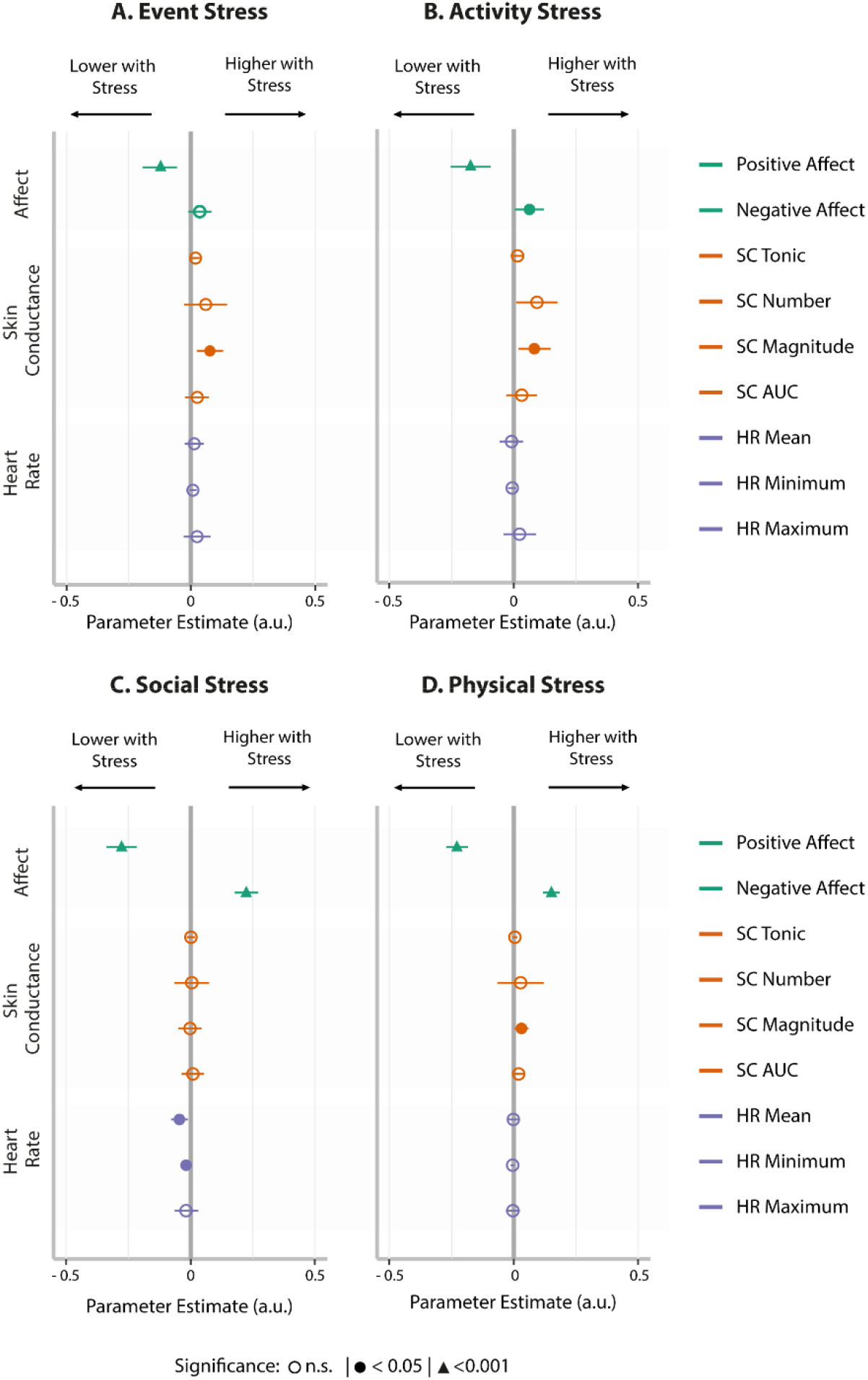
Effect estimates for the associations between moment-to-moment fluctuations in subjective stress and measures of mood and physiology. Subjective stress measures are generally associated with a decrease in positive affect, an increase in negative affect, and increases in some of the measures of physiological arousal. P-values corrected for multiple comparisons using FDR. Error bars represent confidence intervals.

### Positive mood is related to increased arousal and mediates week changes

To investigate whether the observed decreases in physiological arousal during stress weeks could instead be linked to reduced positive affect, we tested the moment-to-moment association between affect and physiological arousal. Increased positive affect was related to increase in the number of skin conductance responses (β=0.08, CI=[0.02,0.06], p_fdr_=0.04), and mean (β=0.01, CI=[0.001,0.02], p_fdr_=0.01), minimum (β=0.01, CI=[0.001,0.02], p_fdr_=0.03), and maximum HR (β=0.01, CI=[0.001,0.02], p_fdr_=0.03, Figure 4, full results in SM Table 4). Thus, in addition to subjective stress, positive affect is also positively associated with momentary physiological arousal.

**Fig 4.**
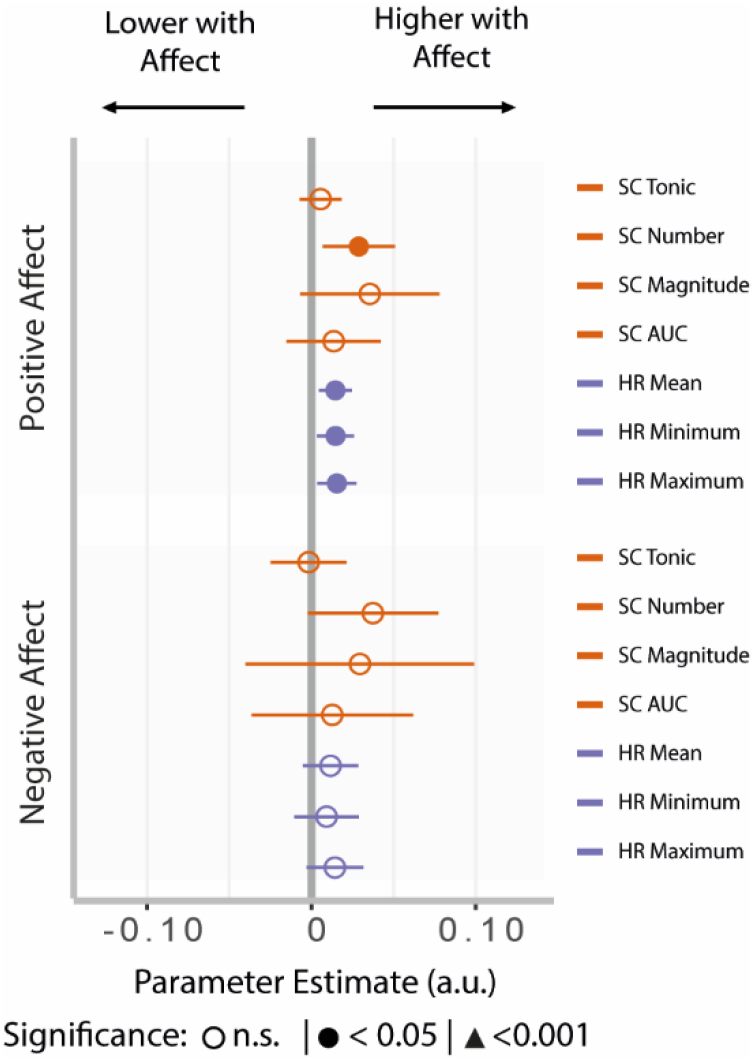
Relationship of momentary affect and physiology. Arousal-related physiological measures (magnitude of skin conductance responses and mean and minimum heart rate) were linked to positive affect, but not to negative affect. P-values are corrected for multiple comparisons using FDR. Error bars represent confidence intervals.

**Fig 5.**
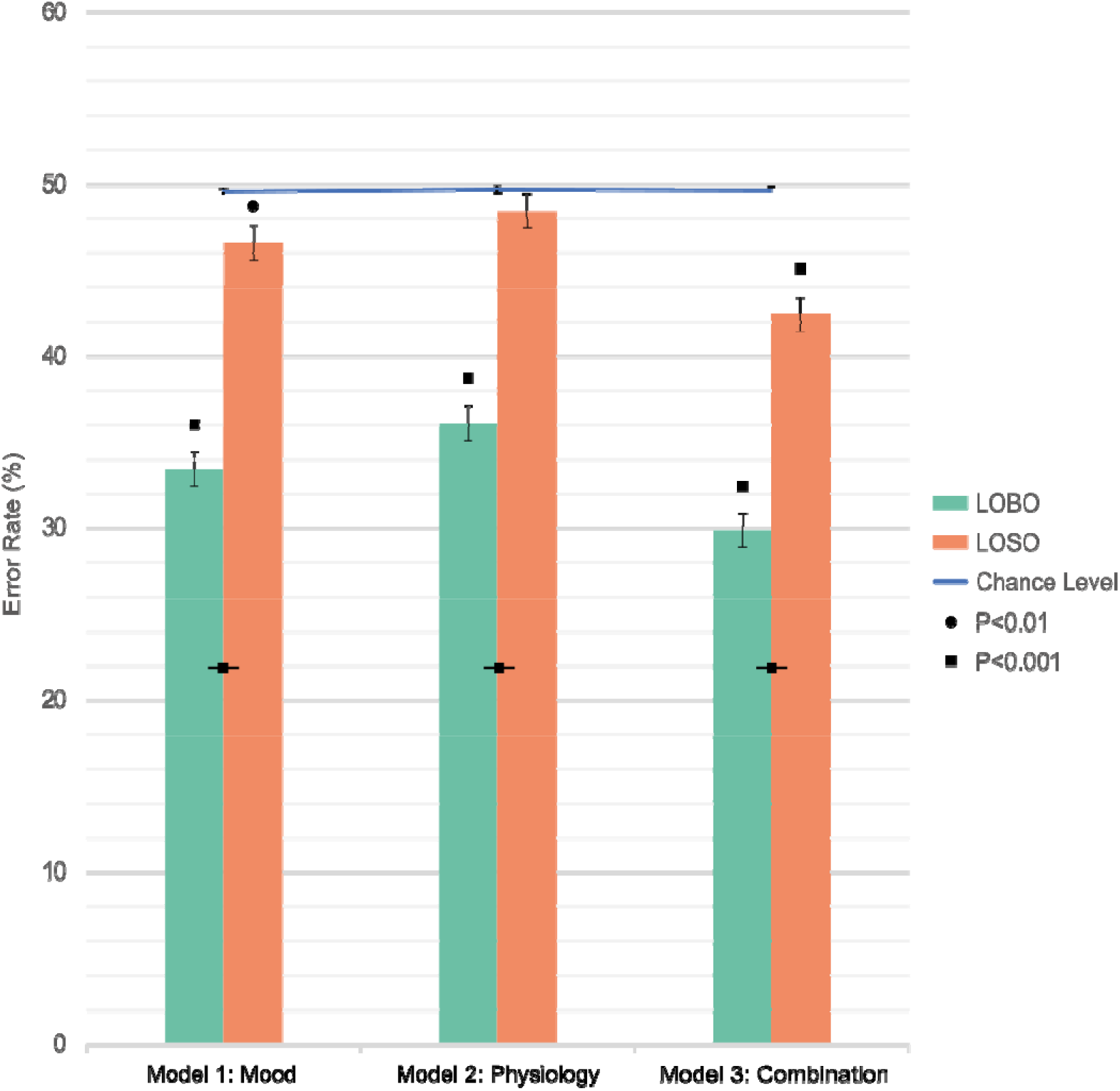
Random-forest classification error estimates. Average error estimates and error bars (representing standard errors of the mean) for each of the random-forest models. Combinations of mood and physiology yield superior classification, and individually trained and tested models (LOBO, Leave-One-Beep-Out) perform better than models trained on group-level data (LOSO, Leave-One-Subject-Out). Chance levels estimated from permutation test and confidence interval are shown in blue. Significance levels between bars indicate between model comparisons, and above bar indicate model comparison to chance levels. P-values corrected for multiple comparisons using FDR.

Next, to confirm that the observed average decrease in physiological arousal observed during the stress weeks is due to the decrease in positive affect, we assessed whether positive affect statistically mediated the effects of week type on reductions in physiological arousal. We specifically focused on the arousal measures linked to subjective stress and positive affect: The number of SC responses and their magnitudes. Results indicated that positive affect significantly mediated the relationship between SC magnitude and week type (7.3%, Mediating Estimate= −0.013, CI=[−0.03, 0.00], p= 0.03) but not fully (Direct Estimate=-0.166, CI=[−0.23, −0.10], p<0.001), indicating potential additional mechanisms are at play. The effect of week type on number of SC responses was not mediated by positive affect.

### Machine-learning classification of beeps using mood and physiology

We next examined to what extent prolonged stress (i.e., stress versus control week) can be classified from individual beeps using machine learning based on affect, physiological arousal, or a combination of both using individualized LOBO models. The mean subject-level error was 33.45% (SD=±2.21) for Model 1 (based on affect), 36.11% (SD=±2.72) for Model 2 (based on physiology), and (M=29.87%, SD=±3.45) for Model 3 (based on affect and physiology combined). Hence, the combined model outperformed the single variable models. All models performed significantly above chance on an individual level for all but one subject (Figure 6, see SM Text 1 for full report). Group-level effects were further tested with paired-samples t-tests with FDR correction comparing the LOBO models to the mean bootstrapped error. Model 1 (Affect, M_diff_=-16.29, t_(80)_=-64.06, p_fdr_<0.001), Model 2 (Physiology, M_diff_=-13.87, t_(78)_=-50.38, p_fdr_<0.001), and Model 3 (Combination, M_diff_=-19.45, t_(78)_=-48.94, p_fdr_<0.001) all performed above chance. Paired-samples t-tests comparing the within-subject error rates between the LOBO models showed that Model 3 (i.e., combined EMA and EPA) outperformed Model 1 using mood alone (M_diff_=3.64, t_(78)_=19.20, p<0.001), which in turn outperformed Model 2 using EPA alone (M_diff_=2.60, t_(78)_=14.65, p<0.001). While overall the EMA mood models performed better, in some subjects the models 1 and 2 had almost equivalent performance.

### Individualized models offer better predictions than group-based models

We next investigated the generalizability of these models from an individualized approach to a population-level one (using group level classification), through the leave-one-subject-out (LOSO) crossvalidation. M1 using affect (45.85%, SD±9.50), M2 using physiology 48.42%, SD=±8.05), and M3 using the combination (42.44%, SD=±9.00) were again tested against their bootstrapped counterpart.

For some individual subjects LOSO models performed significantly above chance level (Model 1-Affect n=45(54.1%), Model 2-Physiology n=30(37.9%), and Model 3-Combination n=55(69.6%)) in classifying week type (SM Text 1). Group level analysis using a paired sample t-test showed that only model 1 (Affect, M_diff_=-3.53, t_(80)_=-3.59, p_fdr_<0.01), and model 3 (Combination, M_diff_=-6.55, t_(78)_=-6.81, p_fdr_<0.001) performed better than chance. Model 2 did not perform above chance (Physiology, M_diff_=-1.34, t_(78)_=-1.54, p_fdr_=1.000).

We additionally directly compared the classification errors between the individualized and group models for each subject using a paired-sample t-test with FDR correction. All individualized LOBO models performed better than the group level LOSO models. LOSO Model 1 (Mood) performed significantly worse than the equivalent LOBO model (M_diff_=11.43, t_(80)_=12.17, p_fdr_<0.001). LOSO Model 2 (physiology) was also significantly worse than the LOBO counterpart (M_diff_=8.11, t_(78)_=8.61, p_fdr_<0.001). LOSO Model 3 (combination) similarly performed worse than the LOBO model counterpart (M_diff_=12.61, t_(78)_=11.74, P_fdr_<0.001). In sum, individual models vastly outperformed group-based models.

## Discussion

This study investigated physiological and psychological responses to ecological stressors in daily life (i.e., exam weeks in students) to determine the usability of passive monitoring technologies for detecting prolonged stress. We employed EMA and EPA to track subjective stress, mood, and arousal-related physiology. Our findings confirmed an overall increase in subjective stress during exam weeks. As hypothesized during the stress week, negative affect increased, and positive affect decreased. Contrary to what was expected, lower skin conductance and heart rate arousal measures were recorded during the stress week. At a beep-to-beep time scale, increased subjective stress was associated with increased negative affect, decreased positive affect, and increased skin conductance responses. Interestingly, positive affect was also associated with increased skin conductance responses, and partially mediated the between-week differences in skin conductance we found. Thus, observed decreases in physiological arousal measures were (at least partially) due to a reduction in positive affect. Using a machine-learning approach, we showed that the combination of individual mood and physiology was best able to detect whether individual beeps stemmed from stress or control weeks. We conclude that passive monitoring with wearable biosensors can detect prolonged stress, highlighting the importance of mood measures to dissociate positive and negative arousal.

In line with previous work, the stress week resulted in increases in self-reported stress, validating our paradigm(Brodersen & Lorenz, 2020). We observed expected changes in mood, with increased negative and decreased positive affect. However, arousal measures were surprisingly reduced during the stress week. The observed overall decrease in physiological arousal during stress weeks appears at odds with the positive association between moment-to-moment subjective stress and increased arousal in our analysis and previous works(Healey & Picard, 2005; Pereira et al., 2017; Smets et al., 2018). This finding reveals a dissociation between prolonged and acute stress. While prolonged stress leads to increased moment-to-moment peaks in self-reported acute stress, it also results more generally in decreased positive affect and decreased overall average arousal. Our results suggest that reduced arousal may be linked to reduced positive affect seen in the stress weeks (irrespective of peaks in subjective stress). Our mediation analysis corroborates this mechanistic link, confirming that positive affect (partially) mediates the effect of week type on reduced arousal. While this may seem counterintuitive, skin conductance and heart rate measures are known to respond to both positive and negative events, showing that physiological arousal is not valence specific(Löw et al., 2008; van Halem et al., 2020). Within a theoretical framework of affect dynamics, these findings also align with the circumplex theory of emotion and valence, linking the two on a grid-like schema of valence and arousal(Posner et al., 2005). Thus, the net effect of prolonged stress exposure stems from a reduction in overall arousal driven by reductions in positive mood that persist outside of peak moments of acute stress.

We subsequently tested the ability of machine-learning models to classify individual beeps as stemming from stress or control weeks with physiology, mood, or both combined. Physiology models could classify beeps almost as well as mood models (3.85% difference on average). However, and more importantly, combination models showed the highest accuracy. Hence, the addition of mood questions to physiological arousal provides valuable information to prolonged stress detection. This converges with the mixed models and mediation results: accounting for valence through mood is necessary to distinguish stress-induced from positive affect-induced arousal. Our findings provide a mechanistic explanation for why previous studies using skin conductance trigger based EMA to detect stress captured positive arousal instead(van Halem et al., 2020). In addition to demonstrating that affect and arousal offer better than chance classification levels, we also show that they achieve higher accuracy in classifying week type than classification based on a median split across explicit subjective stress measures. Assessing mood and physiological arousal may offer a more nuanced measure of stress states that is not dependent on activities or events that occurred since the previous beep. This approach is also common in laboratory research on stress, where mood questionnaires and physiological arousal measures are often used to quantify stress (for example, see (Zhang et al., 2022). In sum, combining a wearable biosensor with minimally invasive mood assessment might offer the best approach to detect stress in both healthy and clinical populations, offering a more feasible approach than full EMA batteries.

Besides demonstrating the utility of physiological monitoring, our results highlight the importance of individualized approaches in stress detection. Classification models trained and tested on individuals’ own data (LOBO) performed significantly better than those trained on group data (LOSO). Our individualized approach offers drastic improvements to classification of stress states, in comparison with group approaches(Smets et al., 2018). Intuitively, the same experience can generate different physiological and psychological responses in different individuals based on a multitude of factors such as sex, appraisal, or clinical traits. For example, patients with anxiety may display a very different physiological response to stress than those with depression (hyper vs hypo activation)(de Looff et al., 2019). This is a key strength of the current approach, fully aligned with recent developments in personalized psychiatry: Individualized models allow for greater prediction accuracy than a one-size-fits all approach.

Worth noting is that classification accuracy of our ML models was relatively low in our study compared to many other ML studies. Importantly, this stems from the ecological design: our models did not classify weeks, but rather individual beeps within the weeks (approximately 70% of beeps were correctly assigned to the weeks in the best models). Through the median split of our data based on self-reported stress, we clearly demonstrate that even during a stress week, participants are not stressed 100% of the (measured) beeps. This was an intentional design choice: the goal was to test the ability of physiology and mood measures combined to detect momentary stress states during heightened periods of stress, which may be required, e.g., for detecting warning signals. Furthermore, the accuracy achieved with our real-life models is also on par with more recent laboratory studies classifying affect from wearables and infrared cameras(Filippini et al., 2022). Hence, classification accuracy found in our study represents what might occur in the general population during real-life stress periods (including stressful moments in regular weeks, and regular moments in stressful weeks), and supersedes directly asking about stress.

Our current study provides early evidence for the successful detection of prolonged periods of stress in individuals. However, some limitations warrant discussion. The current study was purely cross-sectional, but more research (some of which is currently underway, see Healthy Brain Consortium et al., 2021) will be needed to extend these results into prospective stress detection algorithms. Prospective detection of vulnerability is the next step in this line of research, with the aim of identifying early warning signals(Wichers et al., 2020). Such prospective studies are necessary for extending our findings further. It may also be argued that the uncontrolled nature of the study is a detriment to the findings. However, the ecological validity of our study is rather a strength in providing a necessary translation of laboratory measures to a real-life setting (Janssen et al., 2021). Additionally, we controlled for several potential confounds, such as those differences in behavior across weeks (i.e., alcohol intake, sleep, caffeine, exercise). Finally, and worth noting is that an exam stressor may resemble many real-life stressors and daily hassles such as work deadlines. Yet, these results may not generalize to more severe or traumatic stressful life events. Acute stressful events may lead to very different arousal responses, and future research is needed to address this topic. However, having to rely on the occurrence of such events in a study may prove difficult, and would require longer periods of assessments with the hopes of capturing these types of acutely stressful moments. Indeed, this is an issue that was already face by previous attempts of classifying stress from such devices(Smets et al., 2018).

In conclusion, our study shows that EPA may be used for monitoring stress-related mental health but highlights the importance of affect ratings to dissociate changes in arousal due to stress vs. positive affect. A combination of physiology and mood measures is optimal for detecting prolonged stress, and personalized approaches to modeling these variables are necessary. If successfully implemented at a wider scale, our findings may have implications on disease prevention, potentially helping to reduce the overall disease burden of stress-related disorders through personalized early-warning systems and treatment strategies.

## Acknowledgments

The authors would like to acknowledge Mike van Engelenburg from the technical group for his assistance with setting up the EMA platform, and Annemieke Smeets for her assistance with recruitment.

## Data Availability

Data is not made publicly available due to the sensitive and potentially identifiable natured of the data set. All associated raw data used in this study is available upon request via our institutional repository (data.donders.ru.nl). Data repository also includes all scripts used in preprocessing and statistical analysis, which are also included on the GitHub repository linked in SM text 1.

## Author Contributions

Conceptualization and set-up: RT, EH, MK, FK

Data Collection: RT, NK, BK

Data analysis: RT, EV, EH

Writing RT, EV, EH

Editing: All authors

Funding: EH

