## Supplementary Text for "Using wearable biosensors and ecological momentary assessments for the detection of prolonged stress in real life"

**SM Text 1: Survey, Data, and code access**

All surveys, scripts, and analyses code are provided in the online GitHub directory. The full EMA questionnaires are provided in the original Dutch format with accompanying translations.

Results of the random forest models (Leave-One-Beep LOBO, Leave-One-Subject-Out LOSO, and Bootstrapped model) are presented in the interactive notebook hosted on the same GitHub directory. The interactive notebook also contains the individual subject-level and group-level p-values.

Link to interactive notebook: https://raytut.github.io/DetectingStress

Link to code: https://github.com/raytut/DetectingStress

**SM Text 2: EMA and EPA Data Processing**

EMA surveys consisted of questions regarding subjective stress used for validating our experimental paradigm, and mood questions (positive and negative affect) relating to our subjective outcome measures filled in on a 7-point Likert scale. Questions in the validation set probed four types of stress as follows: i) Event-related stress assessed the most prominent event that occurred in between EMA beeps ii) Activity-related stress questions probed the activity participants were engaged in upon receiving the beep iii) Social-related stress addressed stress that may arise from the social context participants were present in (either being alone, or with someone) iv) Physical-related stress was used as a control measure to account for environmental and physical demands. Mood outcome questions consisted of four items assessing positive mood, and five items assessing negative mood. EMA items on a reversed scale were first inverted. Items for each scale were summed to create a single score for each of the scales (i.e., a single measure for Event, Activity, Social, and Physical Stress). Total item scores were then rescaled, and a subject centered measure was derived. Surveys that were not filled in within the assigned time window were excluded from further analyses. The same was done for outcome measure items relating to positive and negative affect.

EPA Data cleaning was performed using Python (V3.6.1)(van Rossum & Drake, 2011). Additional packages used for preprocessing included NumPy (V1.18.1)(Harris et al., 2020) and Pandas (V1.0.3)(McKinney, 2010). Time stamps for each survey instance were used to classify surveys as belonging to a stress or control week. Ten-minute time windows prior to each survey were selected for the extraction of physiology features acquired from the E4. Pre-processed IBI data were deemed too sparse to offer meaningful temporal domain analysis, with an average of 27% of IBIs successfully detected in our selected time window. This is within the margins of the manufacturer's signal loss estimates in daily use. We instead selected average heart rate features from the resulting processed files from Empatica. The devices use a strict proprietary detection algorithm in the detection of IBIs, so these files can be used with minimal processing to derive global heart rate features. These features included the mean, minimum, and maximum heart rate. Raw skin conductance was processed for offline use with the PyPhysio package (V2.1)(Bizzego et al., 2019). A minimum threshold of 0.01 µsiemens was set for the skin conductance levels deemed of acceptable quality based on previous recommendations of a threshold between 0.01-0.05 µsiemens(Boucsein et al., 2012). Data was first despiked to remove artifacts due to sudden hand motions using standard settings in the library. Data was then denoised to remove remaining artifacts through windowed filtering of changes in the signal greater than 0.02 µsiemens between subsequent samples. Additionally, an Elliptic filter with cut-off frequency set between 0.8 and 1.1 was applied to the data. Skin conductance data were subsequently de-convolved using a Bateman impulse response function into phasic and tonic components from which specific features were extracted (mean tonic activity, and magnitude, area under the curve, and the number of phasic responses). The raw temperature measures were used to calculate the mean skin temperature, as well as the slope as a function of change in skin temperature within the acquired time window. Two participants had a watch with faulty temperature sensors. These measures were substituted from the population mean and standard deviation to avoid loss of participants' data due to missing data points in statistical models. The other sensors on this device were tested and no errors were detected in other recordings. Finally, the root mean squared displacement in each time window was calculated from the accelerometer data. The extracted features were collected into a single data frame used for statistical analysis.

**SM Text 3: Statistical Model Fitting**

A maximal fitting approach was used in constructing all our statistical models to reduce Type-I errors in which random slopes and intercepts were estimated for all fixed effects of interest(Barr et al., 2013). In all models, subject was modeled as a random effect, and the variable of interest modeled with a random slope and intercept. Covariates that were factors with 6 or fewer levels were modeled as fixed effects without random slopes. Continuous covariates and those with more levels were modeled as fixed effects with random slopes and fixed intercepts. Covariates were selected to control for potential population differences, and behavioral differences that may arise from being in an exam period, which can be divided into subject level, day level, and beep level covariates. Subject-level covariates modeled as fixed effects included sex, study program, order of the weeks (i.e., stress or control week first). Day level covariates included the days relative to start (i.e., day 1, 2, 3, etc.…), beep number, sleep duration, and the previous night's alcohol consumption. Beep level covariates modeled included hunger, caffeine intake, exercise, and sexual activity. Additionally, ambient temperature and movement were modeled for the EPA models. The same set of covariates was used in all reported models in this paper. For the models assessing moment-to-moment relationships between subjective stress measures and outcomes - due to high correlations between activity stress and both event ($r=0.42$, $p<0.001$) and social stress ($r=0.34$, $p<0.001$) - interaction terms were also modeled for these two variables. Modeling the correlation regresses the change in one variable in relation to the other, thus providing a method for controlling for these effects. For models looking at the relationship between mood and physiology, interaction terms were modeled for positive and negative affect due to high correlations ($r=-0.57$, $P<0.001$). Model fits were checked, and model families were adjusted to achieve optimal fit based on Akaike Information Criterion (AIC) and residual normality. Multicollinearity was checked on base models without the interaction terms. All VIF's of all models were below five.

**SM Table 1**

Shows the results of mixed models looking at the effects of week type on subjective stress ratings. Subject was modeled as a random effect, and covariates as fixed effects and random slopes with fixed intercepts. Random intercepts were added for the fixed effect of interest. Additional covariates modeled include beep and day as factors with interaction terms.

**SM Table 2**

Shows the results of mixed models looking at the effects of week type outcome measures (i.e., positive, and negative affect, skin conductance, and heart rate). Subject was modeled as a random effect, and covariates as fixed effects and random slopes with fixed intercepts. Random intercepts were added for the fixed effect of interest. Additional covariates modeled include beep and day as factors with interaction terms P-values adjusted for multiple comparisons using FDR.

**SM Table 3**

Shows the results of mixed models looking at the momentary associations between subjective stress and outcomes. Interaction terms set for variables with high correlations (Activity Stress and Event Stress, and Activity Stress and Social Stress). Subject was modeled as a random effect, and covariates as fixed effects with random slopes and fixed intercepts. Effects of interest were modeled with correlated random slopes and random intercepts were added. P-values adjusted for multiple comparisons using FDR.

**SM Table 4**

Shows the results of mixed models looking at the momentary associations between affect and physiology measures. Skin conductance shown on left, and heart rate on right. Interaction terms were modeled for positive and negative affect.  Subject was modeled as a random effect, and covariates as fixed effects with random slopes and fixed intercepts. Effects of interest were modeled with correlated random slopes and random intercepts were added. P-values adjusted for multiple comparisons using FDR.
